# An in-silico investigation of the effect of changing cycling crank power and cadence on muscle energetics and active muscle volume

**DOI:** 10.1101/2023.09.18.558368

**Authors:** Cristian D. Riveros-Matthey, Timothy J. Carroll, Mark J. Connick, Glen A. Lichtwark

**Affiliations:** School of Human Movement and Nutrition Sciences University of Queensland Brisbane, QLD, Australia; Faculty of Health. School of Exercise & Nutrition Sciences Queensland University of Technology Brisbane, QLD, Australia

**Keywords:** Cycling, Metabolic Power, Biomechanics, Energy, Simulation, Validation

## Abstract

This study used musculoskeletal modelling to explore the relationship between cycling conditions (power output and cadence) and muscle activation and metabolic power. We hypothesized that the cadence that minimized the simulated average active muscle volume would be higher than that which minimized the simulated metabolic power. We validated the simulation by comparing predicted muscle activation and fascicle velocities from select muscles with experimental records of electromyography and ultrasound images. We found strong correlations for averaged muscle activations and moderate to good correlations for fascicle dynamics. These correlations tended to weaken when analyzed at the individual participant level. Our study revealed a curvilinear relationship between average active muscle volume and cadence, with the minimum active volume being aligned to the self-selected cadence. The simulated metabolic power was consistent with previous results and was minimized at lower cadences than that which minimized active muscle volume across power outputs. Whilst there are some limitations to the musculoskeletal modelling approach, the findings suggest that minimizing active muscle volume may be a more important factor than minimizing metabolic power for self-selected cycling cadence preferences. Further research is warranted to explore the potential of an active muscle volume based objective function for control schemes across a wider range of cycling conditions.

## Introduction

Cyclists typically choose to pedal at a rate that is higher than that which minimizes metabolic power (Coast and Welch 1985; Lucia et al. 2002). It is currently unknown why there is a mismatch between the self-selected cadence (SSC) and the cadence which minimizes muscle energy consumption. However, muscle excitation (measured using electromyography - EMG) measured per cycle has been reported to align approximately with cycling cadence preferences under certain conditions. For example, the average EMG per cycle across different muscles shows a ‘U’ shaped relationship with cadence at fixed power outputs, and the local minimum shifts between 80 and 95rpm as the crank power requirements increase (MacIntosh, Neptune, and Horton 2000; A. P. Marsh and Martin 1995; Takaishi et al. 1996; Neptune, Kautz, and Hull 1997). These local minima roughly correspond to the SSC in submaximal conditions (i.e. ∼94rpm at 10% of the Pmax and ∼106rpm at 30% of the Pmax) (Riveros-Matthey et al. 2023). Although these similarities suggest that an objective function to minimize muscle excitation might apply during cycling, the evidence is limited by technical issues such as the potential for EMG crosstalk, cancellation artefact and an inability to capture deep muscles. Further, EMG metrics are difficult to relate to energy consumption because they do not account for differences in the size of muscles (Sartori, Farina, and Lloyd 2014; Wickiewicz et al. 1984), nor the energy associated with performing mechanical work (Barclay and Curtin 2023; Fenn 1923).

The active muscle volume is an estimated variable that has been linked to the metabolic cost of locomotion in many terrestrial mammals (Roberts et al. 1998; Beck et al. 2019; Biewener et al. 2004). The active muscle volume represents how much muscle tissue is activated to generate required force and therefore provides a metric that reflects to the number of muscle fibers activated, which is one of the key parameters contributing to energy consumption (Beck et al. 2019; Umberger 2010). As such, active muscle volume might be a key factor that the nervous system optimizes during movement. Importantly, unlike activation alone, active muscle volume accounts for the fact that some muscles consume more energy than others because of differences in accounts muscle length and cross-sectional area (Rall 1985; Roberts et al. 1998; Taylor 1994). Presently, the active muscle volume has been typically quantified by combining measures of joint torque with morphological data from human cadavers (Kipp, Grabowski, and Kram 2018; Biewener et al. 2004). However, this approach may ignore muscle dynamics that constrain the force-generating capacity of the muscle (e.g., muscle length and velocity) and contribute to activation requirements.

Musculoskeletal simulations are feasible alternatives to estimate muscle function during human behavior (Delp et al. 1990; Arnold et al. 2013), including cycling (Umberger, Gerritsen, and Martin 2006; Lai, Arnold, and Wakeling 2017; Lai et al. 2021). Modelling approaches enable the estimation of muscle forces that would otherwise be impossible to determine experimentally in tasks such as cycling. By parameterizing the force-generation properties of muscles (Hill 1938; Millard et al. 2013), a variety of forward and tracking solutions (Neptune and Hull 1998; 1999; Crowninshield and Brand 1981; Buchanan et al. 2004) have been developed to solve the muscle redundancy problem, typically using some optimization criterion to characterize the muscle mechanical behavior required to achieve a given movement. As such, muscle activation and muscle force, length and velocity of muscle fibers can be determined (Arnold et al. 2013; Millard et al. 2013; Anderson and Pandy 2001), along with estimation of local muscle energy consumption based on these outputs (Umberger 2010; Umberger, Gerritsen, and Martin 2006). However, such models are often purpose built and require validation under a range of conditions (Park, Caldwell, and Umberger 2022), including the sparse mechanical conditions achievable during cycling.

The purpose of this study was to validate a direct collocation (prescribed) method to characterize muscle dynamics during cycling. Specifically, because the main drivers of energy consumption are muscle activation and active shortening requirements (length change and velocity), we compared simulated muscle excitation and fascicle shortening to experimental EMG and ultrasound data to validate the muscle outputs from the simulation. We then used the model to simulate active muscle volume and muscle metabolic cost across a wide range of cadence and external power requirements. We hypothesized that the model would predict that average active muscle volume (summed across muscles) would exhibit a U-shaped behavior across cadences, with the minimum close to the self-selected cadence, and that this minimum would shift to higher cadences as power demands increased. Furthermore, we hypothesized that the cadence to minimize average active muscle volume would be higher than that which minimized metabolic cost under all power output conditions, consistent with the mismatch between the self-selected cadence and the cadence that optimizes metabolic efficiency (Brisswalter et al. 2000; Alejandro Lucia, Jesus Hoyos 2000; Anthony P. Marsh and Martin 1997; Brennan et al. 2019).

## Material and methods

### Experimental data

The experimental data were extracted from Riveros-Matthey et al. (Riveros-Matthey et al. 2023); the procedures are briefly detailed below. The Human Research Ethics Committees (HRECs) of the University of Queensland approved all procedures and all participants provided their informed, written consent. The study considered twelve level 3-4 cyclists (mean and standard deviation, age= 27 ± 9.9 yr., mass= 70 ± 7.9 kg, height= 1.76 ± 0.054 m) who rode 20sec cycling bouts in an ergometer (Excalibur Sport, Lode BV, Groningen, The 112 Netherlands) under 10%, 30% and 50% of the Pmax at 60,70, 80, 90, 100, 110, 120 and the SSC. Kinematic and kinetic data were measured using 3D motion capture (Oqus, Qualisys, AB, Sweden; Qualisys Track Manager) and force-instrumented pedals (Axis, SWIFT Performance, Brisbane, Australia). The vastus lateralis (VL) shortening velocities were measured using two ultrasound transducers (LV7.5/60/96Z, TELEMED, Vilnius, Lithuania), held in series via a 3D printed frame. EMG signals were recorded wirelessly from gluteus maximus (GMAX), vastus lateralis (VL), rectus femoris (RF), biceps femoris (BF) and semitendinosus (ST), tibialis anterior (TA), soleus (SOL) and gastrocnemius medialis (GM) (Myon 320 system; Myon AG, Baar, Switzerland; 2000 Hz). To run the study simulation, all the experimental data were extracted from the last 5s window recording, considering only the second full crank cycle (the crank cycle is 0 at top-dead-center).

### Musculoskeletal model and simulations

We used a musculoskeletal model developed in OpenSim software (Delp et al. 2007; Rajagopal et al. 2016) and adjusted by Lai (Lai, Arnold, and Wakeling 2017). This model represented the human right lower limb, including 10 segments and 26 degrees of freedom. The model was intentionally developed to increase hip and knee flexion, characterized by tasks such as cycling (https://simtk.org/projects/model-high-flex). Mass and inertia of the torso and upper limbs were cradled to the pelvis. The musculoskeletal model was driven by 40 massless Hill-type muscle-tendon unit (MTU) actuators. Each MTU represented a contractile element attached to a series elastic element (Millard et al. 2013). In this, the contractile element was modelled including active force-length and force-velocity, and passive force-length properties, which were normalized to maximum isometric muscle force, optimal fiber length, and maximum muscle contraction velocity (Zajac 1989).

Using the OpenSim *Scaling tool* (Delp et al. 2007), an anthropometrically scaled model (Figure 1) was generated based on marker data for each participant (MEAN and SEM age= 27 ± 9.9 yr., mass= 70 ± 7.9 kg, height= 1.76 ± 0.054m), with maximum marker errors less than 2cm (Hicks et al. 2015). Joint angles during each cycling task were then calculated using the OpenSim *Inverse Kinematics tool* (Delp et al. 2007), with a maximum marker error of less than 3cm. The experimental pedal forces, which were treated using a 12Hz low pass filter, were rotated with respect to the ergometer’s reference system, using a matrix rotation algorithm. To reduce the residual forces given by the inconsistencies between the kinematic outputs and the pedal reaction forces, an additional 12Hz low pass filter was applied in the inverse kinematics solution, and both joint coordinates and force inputs were used in the simulation. Before running the simulation, the MTU’s muscle-tendon parameters such as the lengths of optimal fiber and tendon slack were scaled morphometrically using an optimization technique (Modenese et al. 2016), without altering the muscle-tendon dimension of the scaled Lai model (Figure 1). An energetic probe was included in the model (Umberger 2010; Umberger, Gerritsen, and Martin 2006) to estimate energetic demand (metabolic rate) in each condition. The muscle energetics model considered the effect of muscle mass, the ratio of muscle slow-twitch fibers, and the muscle fiber velocity.

**Figure 1:**
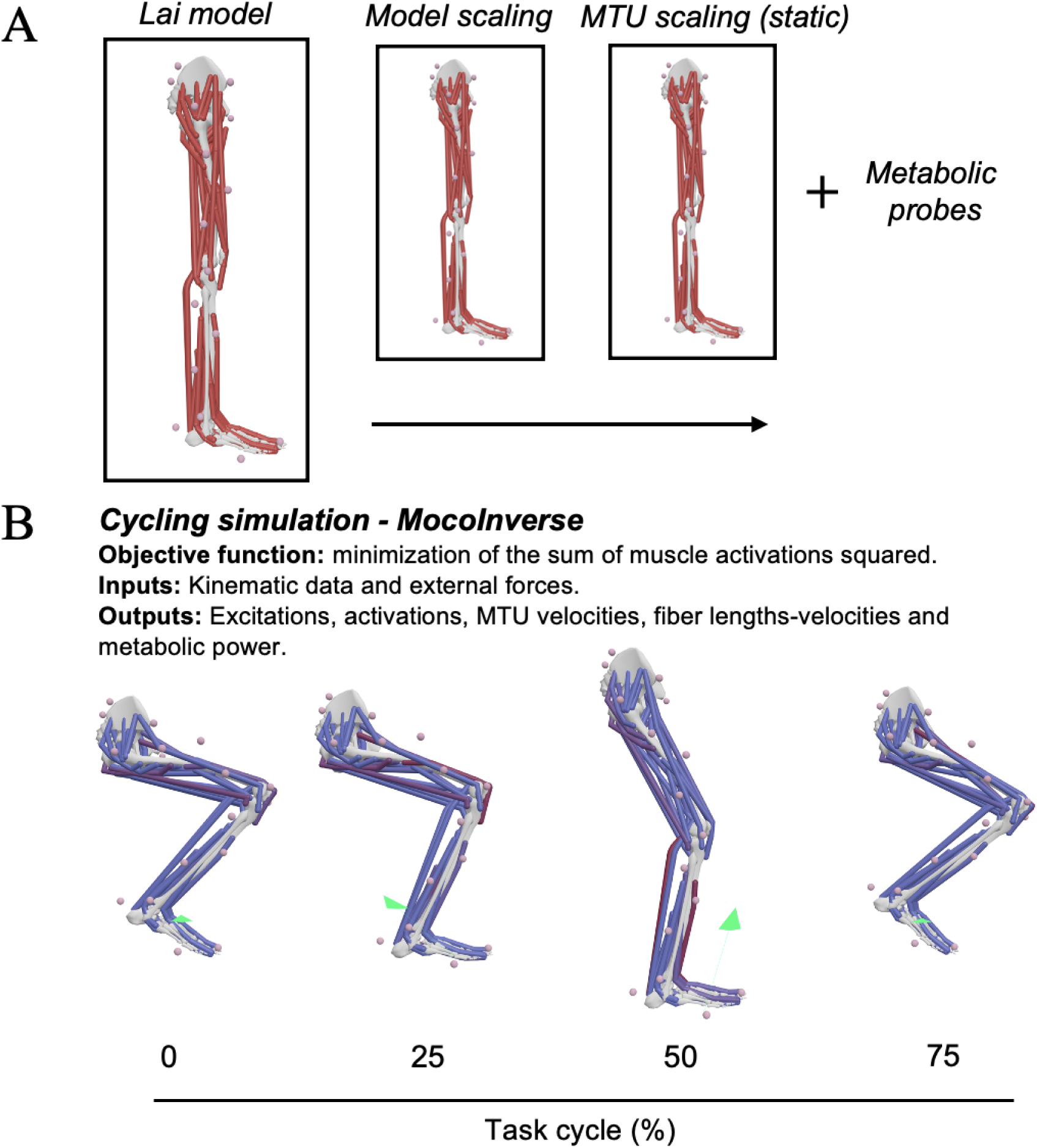
Summary of the prescribed simulation procedure: (A) Musculoskeletal model by Lai (2017) with 40 actuators scaled according to the characteristics of each participant and the muscle-tendon parameters adjusted through the optimization technique (Modenese et al. 2016). (B) Cycle of the simulation where the task cycle (%) was considered as 0 at the top dead center. The green arrow represents the external forces.

Simulations were implemented using OpenSim *MocoInverse*, which solves the muscular redundancy problem using a direct collocation technique, prescribing the motion exactly (Dembia et al. 2019). The scaled model, the inverse kinematic solution (joint angles), and the pedal forces from the experimental design were the inputs. The minimization of the sum of muscle activations squared was the cost function used to solve the redundancy problem. Given the fast motion of the task, a mesh interval of 0.01 sec was used during the simulation. In order to ensure that the controls in the simulation achieve greater smoothness and that the cost function is minimized, a convergence and constraint tolerance of 0.0001 was used. After the simulation, the OpenSim Muscle Analysis tool was used to determine MTU outputs (i.e., MTU force, MTU velocities, fiber lengths, and velocities) and metabolic power demand.

### Data Analysis

#### Model validation

The validation procedure included experimental data from six participants to test the model’s performance. Participants were selected based on their similar anthropometrics extracted from each model, which made them suitable for comparison in the study (mass= 65 ± 1.8 kg, height= 1.75 ± 0.051 m). The experimental EMG-RMS and VL fascicle shortening were compared to those predicted by the simulation technique. The experimental data were processed in the manner described by Riveros-Matthey et al., (Riveros-Matthey et al. 2023); but additional analyses for validation were also carried out. To compare EMG to model excitation patterns, we applied a moving root mean square (RMS) with a window width of 50 ms to both the recorded EMG signal and the simulated excitation patterns for each muscle, which created a smoothed envelope for comparison. The activation and excitation amplitude for each muscle were normalized to the maximum value during the 60rpm cadence at 50% of Pmax. The velocity from the experimental and simulated VL fascicle was computed with respect to the crank angle as the first derivative of their lengths with respect to time. The analysis utilized the group and individual approaches to assess the correspondence between different metrics. At the group level, the average group values (participants) were considered for all conditions (cadences x intensities), resulting in 24 data points. At the individual level, a comparison was conducted by considering individualized values (cadences x intensities) per participant. The inputs for analysis included the averaged muscle EMG RMS activation and simulated excitations, the averaged VL fascicle velocities and peak VL fascicle velocities over a full pedaling cycle. To assess the similarities between experimental and simulated metrics, we used a coefficient of determination (*R*^2^). Linear regression was performed for the group-level analysis, while linear mixed effect models were computed for the individual analysis, adding participants as a random effect.

#### Average active muscle volume

The selected MTUs for analysis were the GMAX, VL, RF, BF, ST, TA, SOL, and GM. The time-series data for muscle mechanics were processed per each cadence condition and crank power requirement considering the average of a pedaling cycle where the crank angle is 0 at the top dead center. We first calculated the volume of each muscle (*V*_*tot*_) using the muscle model parameters:

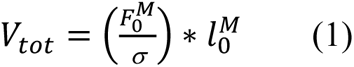

where 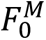 is the maximum isometric force, σ is the isometric force per unit of cross-sectional area of active fibers (assumed to be 20 N *cm*^−2^) (Perry et al. 1988), and 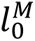 is the optimal fiber length of the muscle. We then calculated the active muscle volume (*V*_*act*_) at each time instant for each muscle:

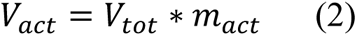

where *m*_*act*_ is the muscle activation level.

To represent the average active muscle volume across all muscles of the lower limb, the average active muscle volume across time (*V*_*sum*_) was computed by averaging the active muscle volume per MTU-actuator across the entire pedal cycle 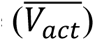 and summing across muscles:

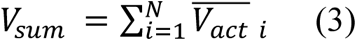

where *i* is the muscle number and *N* is the number of muscles measured (N=8).

#### Metabolic expenditure analysis

The total metabolic power across pedal cycle was extracted from Muscle Analysis tool (Umberger probe; Umberger 2010) after running the simulation (Dembia et al. 2019). The metabolic probe calculates the overall metabolic energy rate (power) of a group of muscles in the model during simulation. The algorithm considers the metabolic power of the muscles, which is equal to the heat release plus the rate of work done, based on muscle states (activation, length, velocity) across time. To calculate the net metabolic power (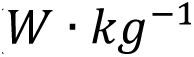), the resting metabolic rate was subtracted from the total instantaneous metabolic power obtained from the probes. Then, the mass-normalized instantaneous metabolic power over the pedal cycle was averaged and divided by the duration of the pedal cycle (Cseke, Uchida, and Doumit 2022). Further, the resulting single-leg average net metabolic power was doubled to account for the energy expended on the lower extremities, based on the assumption that the metabolic energy rate is similar in both legs.

### Statistical analysis

All data presented are group means and standard deviations. To examine the relationship between the total active muscle volume and cadence at each pedal power requirement, the per-muscle and summed active muscle volumes were fit with a second-order polynomial nonlinear regression, which was used to quantify the cadence that minimized active muscle volume per participant. Then, using a linear mixed effect model, the same approach as in the validation (group and individual analysis) was used to assess the correspondence (*R*^2^) between the fixed effect of cadence that minimized active muscle volume (per-muscle and summed form) and the experimentally determined SSC, with participants as a random factor to account for individual variations in slope and y-intercept. Correlations were categorized based on the magnitude of their *R*^2^ values: ≤ 0.35 as weak, 0.36 to 0.69 as moderate, 0.70 to 0.89 as strong, and ≥ 0.90 as very strong. An α level of 0.05 was set for all tests. We also used a second-order polynomial nonlinear regression to examine the relationship between predicted metabolic power and cadence at each pedal power demand. All data were processed in MATLAB (R2022a, MathWorks Inc., USA).

## Results

### Validation results

The model generated muscle excitations that were consistent with the participants’ EMG activations at different power outputs and cadences using the prescribed simulation (Figure 2). The linear mixed effects model revealed that the relationship between averaged EMG activations per condition (power output x cadences=24 conditions) and predicted excitations were relatively strong across muscles (Figure 2). At group level analysis, the muscles with the highest percentage of variation explained by the model were VL (*R*^2^=0.94), BF (*R*^2^=0.92), GMAX (*R*^2^=0.89), GM (*R*^2^=0.84), SOL (*R*^2^=0.83), and ST (*R*^2^=0.80), whilst TA displayed a moderate correspondence (*R*^2^=0.69) (Figure 3). Further, when all individual data was considered with participant as a random factor, the *R*^2^in all muscles decreased, with the RF muscle having the lowest *R*^2^ of 0.37 (Figure 2).

**Figure 2:**
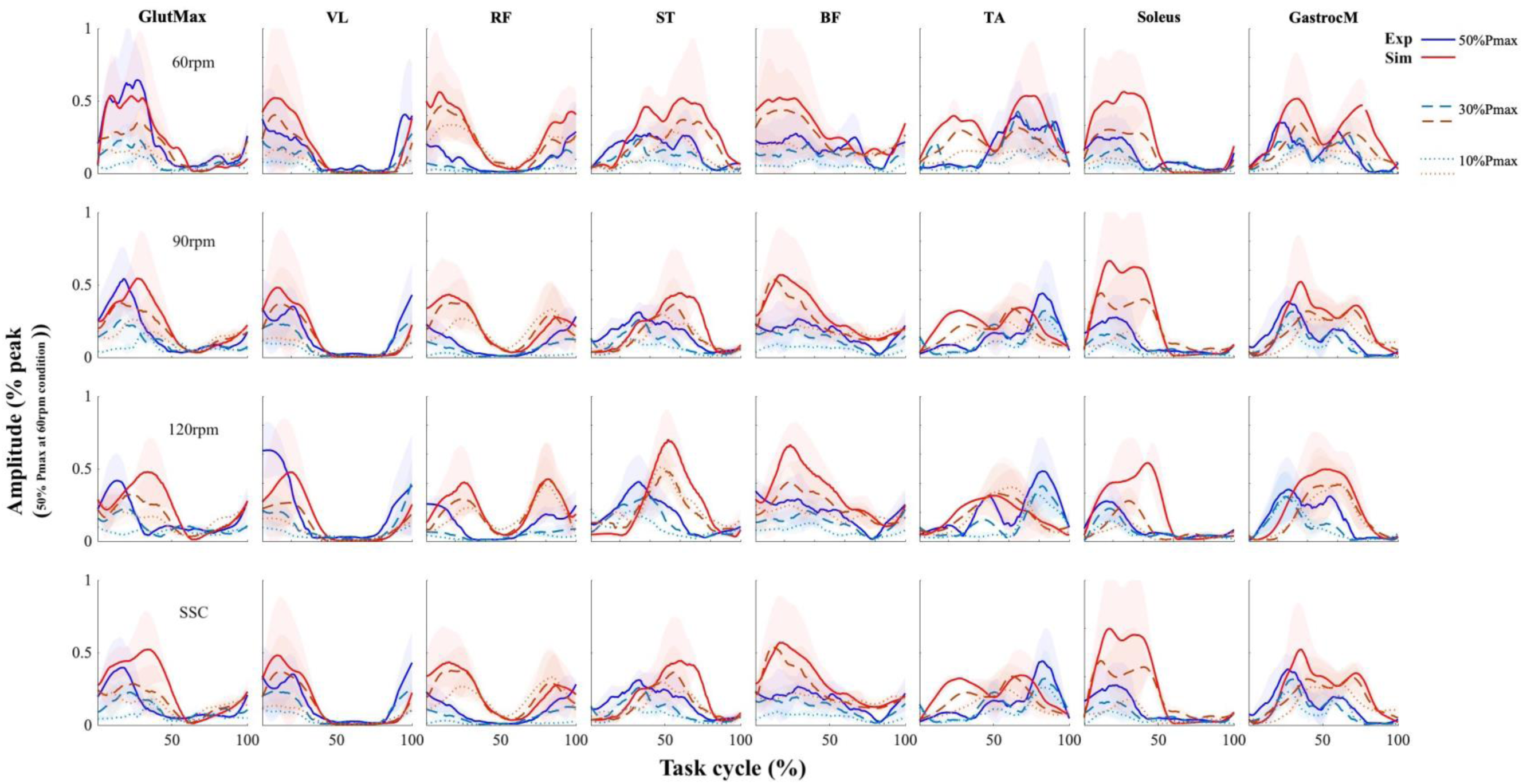
Comparison between experimental EMG activation and simulated excitations of GMAX, VL, RF, ST, BF, TA, SOL and GM muscles across 10%, 30% and 50% of Pmax at 60, 90, 120rpm and SSC. Derived blue colors correspond to experimental muscle activation whereas red ones are from simulated excitations. Straight lines correspond at 50%, dash lines at 30% and dotted lines at 10% of the Pmax. The task cycle (%) is 0 at top-dead-center. The shaded area around lines represents the standard deviation. The 100% amplitude normalization was set considering the peak of the activation and excitation at 50% of the Pmax at 60rpm.

Figure 3 shows the measured and simulated VL fascicle velocities at cadences 60, 70, 80, 90, 100, 110, 120, and SSC rpm at 30% of Pmax to evaluate how well the model replicated the fascicle dynamics across the crank cycle. When the VL fascicle shortening velocities were compared to those predicted across cadences and pedal power demands, the group averaged data yielded an *R*^2^ of 0.85, while the individual (participants as a random effect) analysis yielded a *R*^2^ of 0.66. The peak VL shortening velocities analysis resulted in an *R*^2^ of 0.56 across conditions (power demands x cadences) for the group average and an *R*^2^ of 0.57 for the individual participant effects.

**Figure 3:**
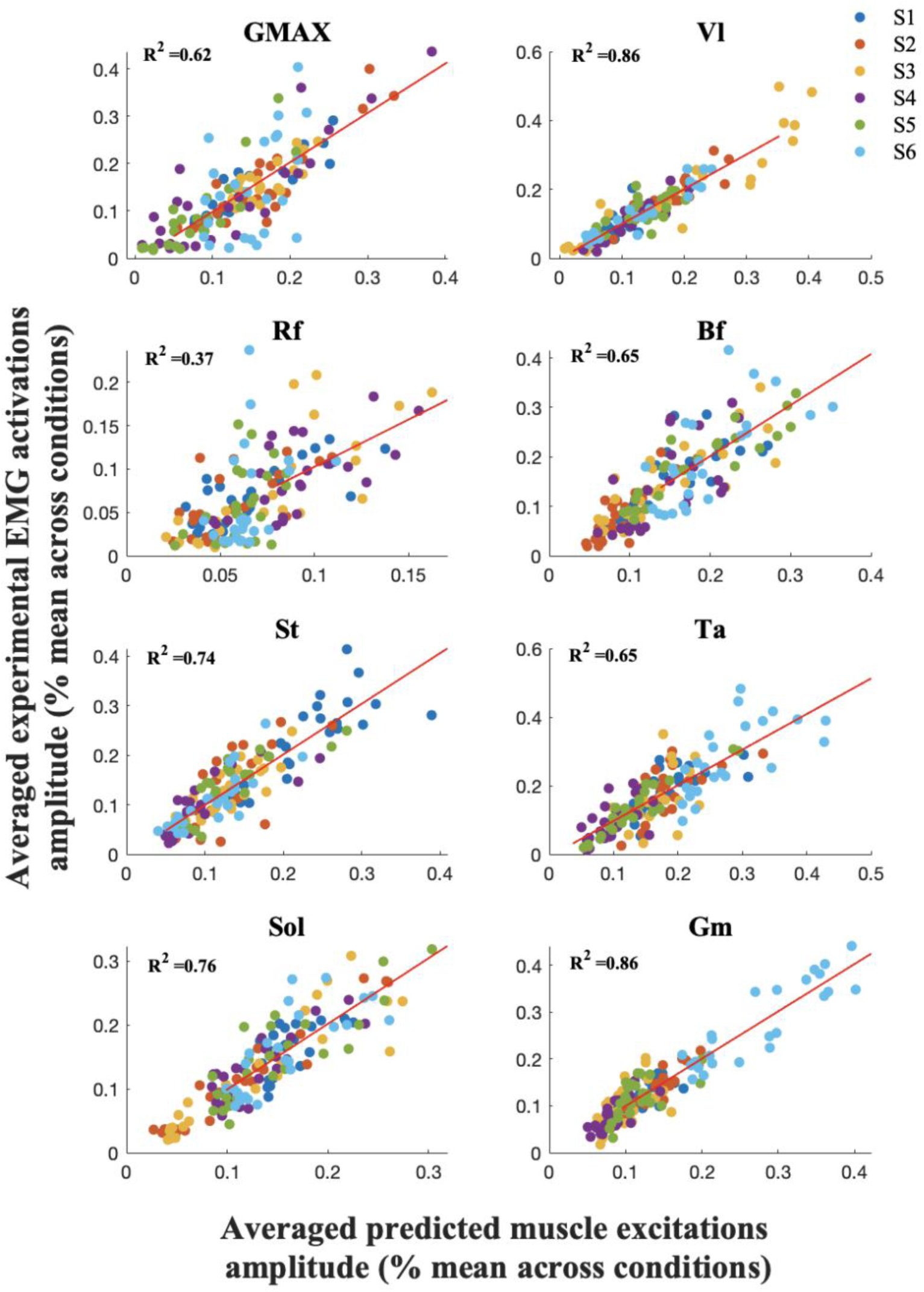
Linear mixed-effect regression model (participants were included as random variables) between averaged experimental EMG activations and predicted excitations across cadences and power outputs (24 conditions) per muscle. Participants are shown as colored dots, and the fit for each participant is shown as colored dashed lines.

**Figure 4:**
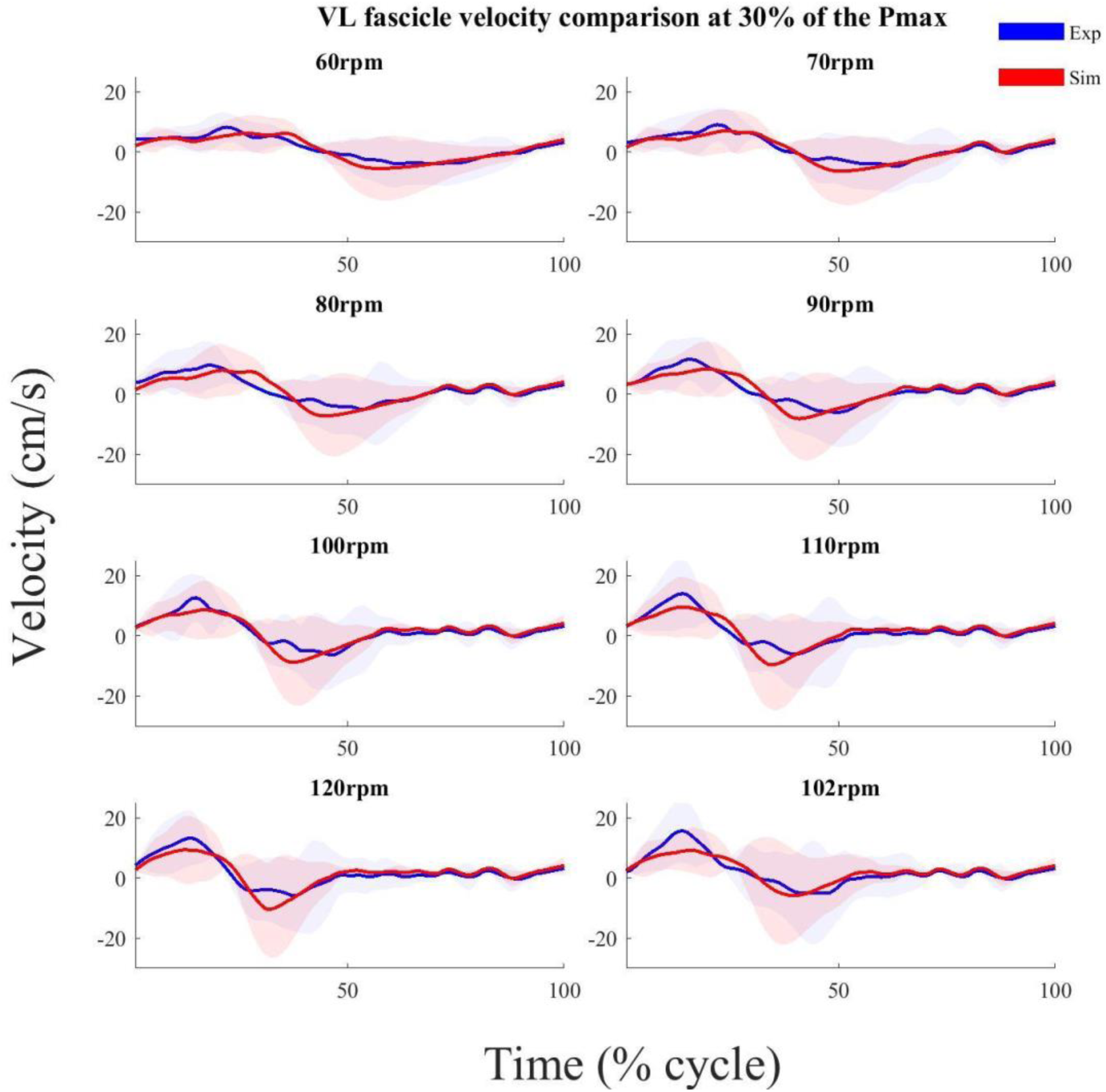
Averaged VL fascicle velocities per cycle at 60, 70, 80, 90, 100, 110, 120rpm and SSC at 30% of the Pmax. The self-selected cadence (SSC) is represented by 102rpm. The shaded area around lines represents standard deviation.

### Active muscle volume and metabolic power results

A curvilinear relationship between average active muscle volume and cadence was observed across muscles (Figure 5), with increasing active muscle volume required for increasing power demands. On average, a strong correspondence (*R*^2^=0.83) was observed across muscles at the group level. The linear mixed-effect model revealed a moderate association between the cadence that minimized average active volume and the SSC across different muscles at the individual level, with an average *R*^2^ of 0.68 across muscles. Notably, a strong correlation was found for the GM (*R*^2^=0.86) and VL (*R*^2^ = 0.86), while the RF exhibited a weaker *R*^2^ of 0.37.

**Figure 5:**
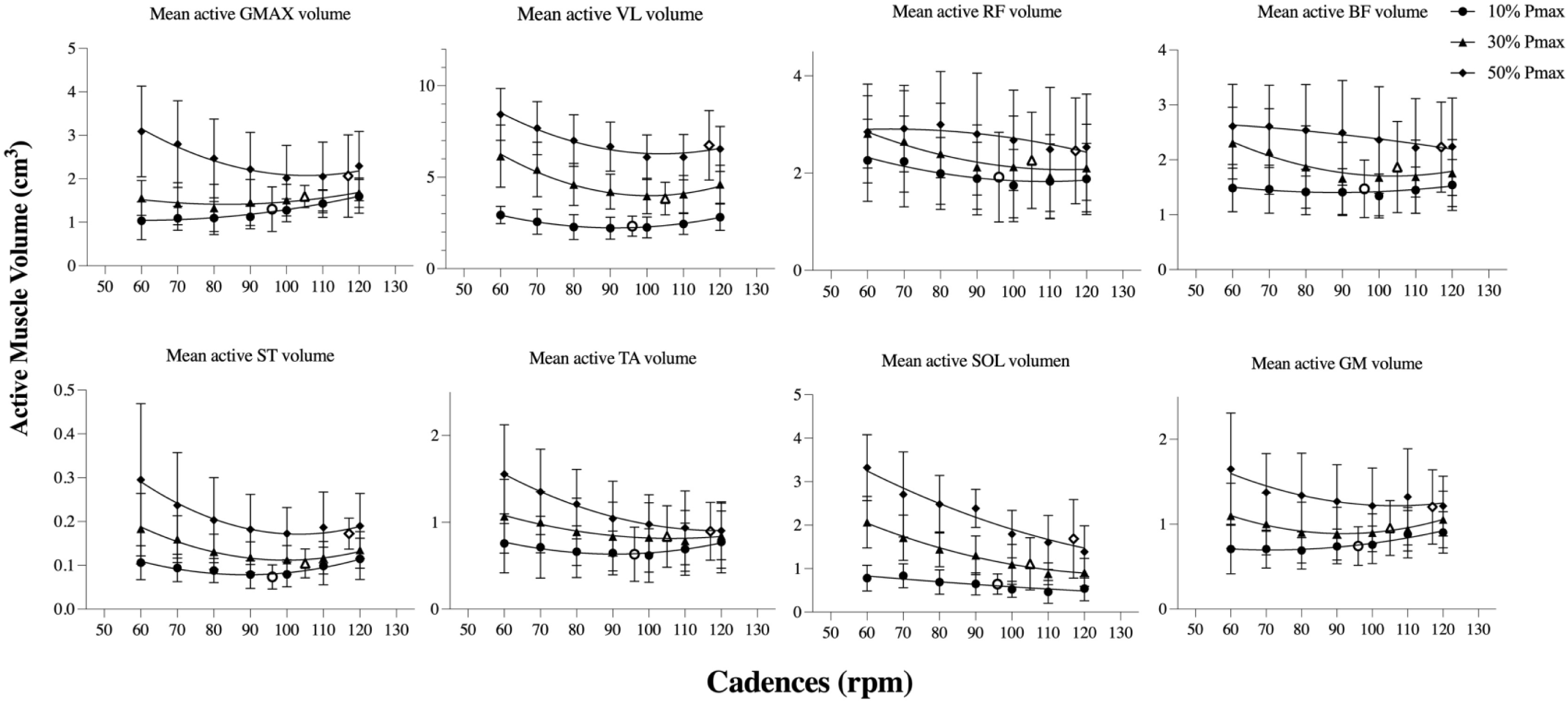
Mean active volume of GMAX, VL, RF, BF, ST, TA, SOL, and GM at 60, 70, 80, 90, 100, 110, 120rpm and SSC at 10% of the Pmax (a highlighted black dot), 30% of the Pmax (a highlighted black triangle), and 50% of the Pmax (a highlighted black diamond). Data shown are means ± SD. Lines represent a second-order polynomial nonlinear regression. The SSC is 97rpm at 10% of the Pmax (empty circle), 106rpm at 30% of the Pmax (empty triangle), and 116rpm at 50% of the Pmax (empty diamond).

A curvilinear relationship was also found between the summed average active muscle and cadence, and the minimum of this relationship shifted to higher cadences as external power demands increased (Figure 6A). While the group analysis displayed a very strong correspondence (*R*^2^=0.99), the individual level analysis showed a moderate correlation between the local minima of summed active muscle volume predicted from the 2^nd^ polynomial regression and SSC across power demand conditions with a *R*^2^ of 0.66 (Figure 6B).

**Figure 6:**
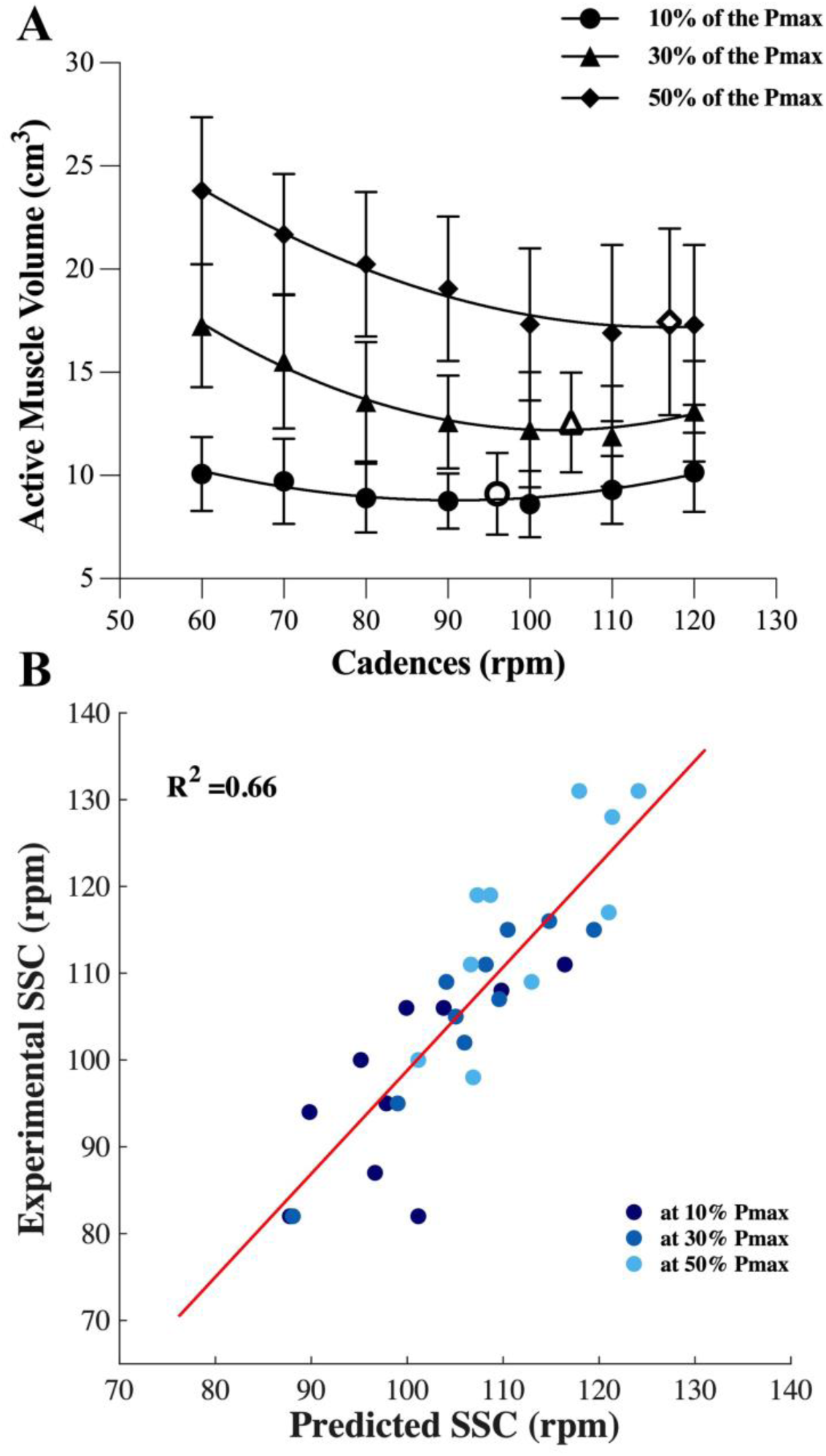
(A) The summed average active muscle volume at 60, 70, 80, 90, 100, 110, 120rpm and SSC at 10% of the Pmax (a highlighted black dot), 30% of the Pmax (a highlighted black triangle), and 50% of the Pmax (a highlighted black diamond). Data shown are means ± SD. Lines represent a second-order polynomial nonlinear regression. The SSC is 97rpm at 10% of the Pmax (empty circle), 106rpm at 30% of the Pmax (empty triangle), and 116rpm at 50% of the Pmax (empty diamond). (B) Linear mixed-effect regression model (participants were included as random variables) between experimental SSC (rpm) and cadences where the summed active volume minima predicted were observed from the nonlinear regression across power demand conditions (10%, 30% and 50% of the Pmax). Participants are represented by dots across power demands (blue dark: at 10% of the Pmax; blue: at 30% of the Pmax; blue light: at 50% of the Pmax) and the fit from the mixed-effect regression model is represented by a red line.

A curvilinear relationship was also found between the predicted metabolic power and cadence, and the minimum of this relationship shifted to higher cadences as the power demands increased (10% of the Pmax: 63 ± 5.4 rpm; 30% of the Pmax: 65 ± 7.1 rpm and 50% of the Pmax: 67 ± 7.5 rpm; Figure 7A, C and E). As has been shown experimentally in numerous investigations (Foss and Hallén 2004; Coast and Welch 1985; Brisswalter et al. 2000), minimum metabolic power occurred at notably lower cadences than the SSC across power (vertical lines in Figure 7A). A comparison between our simulated metabolic power and experimental data obtained across a similar cadence and power range in a previous report (Foss and Hallén 2004) are also shown for qualitative comparison (Figure 7B, D and F).

**Figure 7:**
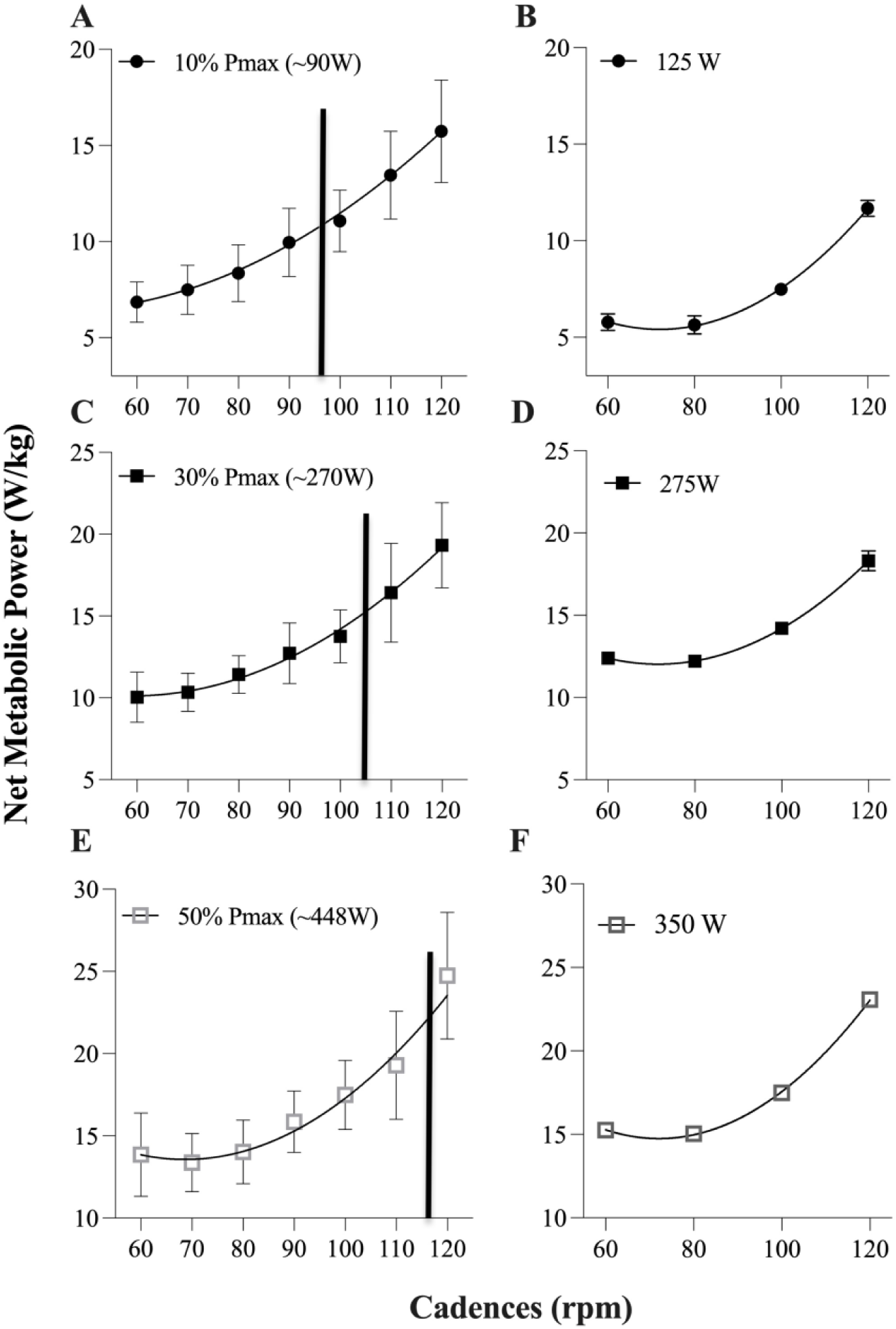
Net metabolic power predicted by the probes at 60, 70, 80, 90, 100, 110, 120rpm and SSC at 10% of the Pmax (A; a highlighted black dots), 30% of the Pmax (C; a highlighted black squares), and 50% of the Pmax (E; a highlighted grey empty squares). Data shown are means ± SD. Lines represent a second-order polynomial nonlinear regression. The SSCs (vertical lines) are 97 rpm, 106 rpm, and 116 rpm at 10%, 30%, and 50% of the Pmax, respectively. Charts B, D, and F show the net experimental metabolic power based on data from Foss and Hallén (2004) at 60, 80, 100, and 120 rpm across power demands (125 W, 275 W, and 350 W), which were initially calculated as *V*02 · min ^−1^. The net energy expenditure (*kj*/min) and an incremental respiratory exchange ratio ((RER); from 0.7 to 1; Péronnet and Massicotte 1991) were considered as cadence and power demands increased in order to convert them to mass-normalized metabolic power (*W/kg*) (Foss and Hallén 2004).

## Discussion

In this study, our goal was to explore the relationship between cycling dynamics, active muscle volume and predicted metabolic cost. The simulation method was capable of reasonable predictions of experimentally measured muscle activations and fiber velocities across a large range of conditions, providing confidence in the estimates of active muscle volumes and metabolic cost. In support of our hypothesis, the summed average active muscle volume exhibited a curvilinear relationship with cadences, demonstrating that the minimum active muscle volume was cradled close to the self-selected cadence, and shifted towards higher cadences as power demands increased. Moreover, although the estimated metabolic cost curve also had a minimum that shifted towards higher cadences as power requirement increased, the metabolic cost minima were much lower than the self-selected cadences. These outcomes indicate that the simulation method can reasonably predict cycling dynamics and energetics. Interestingly, the summed average active muscle volume was not a good predictor of metabolic rate, most likely due to the additional cost of muscle shortening at high rates with increased cadence (Barclay and Curtin 2023; Fenn 1923; Umberger, Gerritsen, and Martin 2006). We also speculate that while energy costs may be lower at lower cadences, the requirement to activate more could contribute to a form of fatigue (peripheral or central) that could limit overall cycling performance over time. Instead, due to the close similarity with muscle activation, which has also been associated as a predictor of SSC, the summed average active muscle volume is more likely acting as a cost of activating muscle for force generation, reducing the demand placed on motor units and muscle fibers, making it a better predictor of cycling behavior than a proxy of energy expenditure. Therefore, it appears that criteria related to muscle activation are more important drivers of behavior during cycling than criteria related more directly to energy consumption rates, given that the minimization of active muscle volume was better aligned to SSC than the minimization of metabolic power.

The study showed reasonable model validity for muscle activation and VL muscle fascicle velocities across different power demands and cadence conditions. Recent studies that used a direct collocation approach during walking (Bianco et al. 2022) and hopping (Jessup, Kelly, and Cresswell 2023) showed a similar correspondence to experimental data. The Umberger metabolic probe also revealed a curvilinear pattern in relationship between cadence and metabolic power for all intensities, which is consistent with previous studies. For example, when comparing our findings to those of Foss and Hallen (2004) - particularly for power demands of 10% of Pmax (90W/125W; Figure 7A and B) versus 50% of Pmax (448W/350W; Figure 7E and F) - we observed a substantial and consistent increase in metabolic consumption across the range of cadences. Whilst further metabolic validation is required, our results provide confidence in the capacity to represent general muscle mechanical and energetic function during cycling using the direct collocation simulation technique.

Our study provides a novel insight into the relationship between pedaling cadence and the active volume of muscle across power demands. We found a clear minimum in the relationship between individual active muscle volumes and cadence for the GMAX, VL, ST, TA and GM muscles (Figure 5). Furthermore, the summed muscle active volume clearly captured a curvilinear relationship across power output conditions (see Figure 6A) with the minimum active muscle volume cadence very well correlated to the self-selected cadence (irrespective of the crank pedal power demands). Our results are in line with studies showing that the muscle activation across muscles is minimized at cadences of ∼90rpm (MacIntosh, Neptune, and Horton 2000; Takaishi et al. 1996), and that this muscle activation minimum is close to self-selected cadences in both simulated (Neptune and Hull 1999) and experimental cycling (Riveros-Matthey et al. 2023). Muscle activation minimization has been attributed to a nervous system strategy to reduce fatigue, whereas a minimization of summed active muscle volume has been referred previously to as a proxy for energy expenditure. Our results suggest that minimizing active muscle volume does not necessarily minimize energy during cycling, and therefore active muscle volume may be a poor proxy of energy use during this largely concentric, power generating task. Given the large number of muscles that minimized their active volume (through minimizing activation) near the self-selected cadence, it is perhaps not surprising that additional weighting for muscle sizes did not alter the relationship between cadence and summed average active muscle volume compared to examining activation in isolation. However, further work is required to test the generalizability of the results across different cycling conditions, including those that may deviate from the optimal and other locomotor conditions.

In line with previous findings (Foss and Hallén 2004; Coast and Welch 1985; Lucia et al. 2002; Figure 7B, D and F), our simulations suggest a mismatch between the cadence which achieves minimum metabolic cost and that which minimizes active muscle volume. These disparities are largely because of additional energy expenditure associated with increased shortening rates observed at higher cadences (Barclay and Curtin 2023; Fenn 1923; Umberger 2010) which can’t be captured by active muscle volume alone. Notably, findings focusing on walking have reported a closer alignment between conditions that minimize energy expenditure and those which minimize muscle activations, relative to cadence (stride rate) (Russell and Apatoczky 2016; Sousa and Tavares 2012). This discrepancy between tasks may be attributed to fact that walking allows storage and return of energy while cycling is a power generating task; the positive work performed by muscles and the higher heat rate observed during higher shortening rates results in increased energy consumption for a given work rate as cadence (and shortening rate) increases. Consequently, a question arises regarding the underlying reasons for the nervous system’s failure to optimize energy-related factors, including the cost of shortening (Fenn 1924) induced by higher cadences (e.g., the SSC) in cycling. One speculative explanation could be that the nervous system aims to avoid excessively low cadences (which are maximally economical) in order to reduce prolonged activation and potential central fatigue (Bailey et al. 2008; Ludyga, Gronwald, and Hottenrott 2016; Hottenrott, Taubert, and Gronwald 2013). While higher cadences may induce greater peripheral fatigue due to the higher energy cost of high contraction velocity (Ludyga, Gronwald, and Hottenrott 2016; Gandevia, Refshauge, and Collins 2002), the local effects of fatigue may be mitigated by reduced intramuscular pressure and increase blood flow due to more frequent muscle relaxation (Hagberg et al. 1981; Takaishi et al. 1998; Abbiss and Laursen 2005). However, further empirical evidence is necessary to corroborate these hypotheses.

### Limitations

The outcomes of this study must be interpreted acknowledging certain limitations. Instead of directly comparing activations and excitations over time, the validation procedure concentrated on analyzing the average values throughout the entire task cycle. This method was chosen due to inconsistencies in the estimation of the electromechanical delay at the onset of simulated excitation (Figure 2), which usually results in a time shift in the output signal when compared to experimental muscle activation over the task cycle (Heine, Manal, and Buchanan 2003). Furthermore, employing the technique of averaging values across the complete task cycle while considering a broad range of conditions, including varying power demands and cadences, serves to mitigate potential sources of data variability, given by time-scale fluctuations, noise, and outliers in individual data points. Altogether, this approach enables the comprehensive capture of a broad spectrum of movements, encompassing fluctuations in speed, alterations in direction, and variations in external loads (Hicks et al. 2015).

We further acknowledge that the utilization of a right lower limb model based on the assumption of symmetry may not fully capture potential asymmetries in muscle activation, joint loading, and hence metabolic power reported in experimental cycling (Carpes, Mota, and Faria 2010). Moreover, the focus of our analysis on local results restricts the ability to generalize these findings to whole-body performance or broader movement tasks. While predicted muscle activations had strong correlations with experimental EMG, this may have been influenced by the cost function used (the sum of muscle activations squared) to solve the redundancy problem in the simulation. Moreover, there was still considerable variation in some muscles (e.g., SOL; Figure 2), particularly when we consider the individual participant analysis. This may be due to assumptions of uniform muscle-tendon properties or other geometrical issues associated with scaling (Hicks et al. 2015). Lastly, although our study incorporated static muscle-tendon unit (MTU) calibration, aimed at improving the representation of muscle-tendon behavior. It is worth noting that a combined static and dynamic MTU calibration approach may offer an enhanced and more realistic depiction of muscle behavior during dynamic activities such as cycling. The integration of dynamic calibration techniques could account for the time-dependent adaptations in muscle properties, yielding a more comprehensive understanding of muscle-tendon behavior in movement scenarios (Pizzolato et al. 2015).

## Conclusion

We showed that a musculoskeletal model provided simulated muscle dynamics with acceptable correspondence to experimental data during a wide variety of cycling task conditions. The simulations showed curvilinear relationships between cadence and active muscle volume, with minima shifting to higher cadences with increased power output, corresponding well with the SSC. This implies that the average volume of muscle (summed across muscles) activated across time is a potential parameter to explain why cyclists do not minimize energy consumption when selecting a cadence. These findings offer a complementary viewpoint to the established role of muscle activation minimization highlighted in existing literature. Simulation across different power demands clearly demonstrates the complex interplay between muscle mechanics and energy demands in lower limb cycling and helps elucidate the factors driving self-selected cadences in cycling tasks.

## Declarations

## Acknowledgements

We acknowledge The University of Queensland for support and provision of resources that were essential for conducting this study.

## Availability of data

Data can be found at https://gitfront.io/r/user-8029681/s8m45dDKLTP4/Simulated-Cycling-Data/

## Authors’ contributions

C.R-M., T.J.C, M.J.C. and G.A.L. conceptualized and designed the experiment. C.R-M. conducted the experiments. C.R-M analyzed data, prepared figures, and drafted manuscript. C.RM., T.J.C., M.J.C. and G.A.L. interpreted results, revised manuscript, and approved final version of manuscript.

